# Optical sectioning for reflection interference microscopy: quantitative imaging at soft interfaces

**DOI:** 10.1101/2024.06.07.598038

**Authors:** Cathie Ventalon, Oksana Kirichuk, Yotam Navon, Yan Chastagnier, Laurent Heux, Ralf P. Richter, Lionel Bureau, Delphine Débarre

## Abstract

Reflection Interference Contrast Microscopy (RICM, also known as interference reflection microscopy) and related techniques have become of wide interest to the biophysical, soft matter and biochemistry communities owing to their exquisite sensitivity for characterising thin films or individual nanoscopic objects adsorbed onto surfaces, or for monitoring cell-substrate interactions. Over the recent years, striking progresses have been made to improve the sensitivity and the quantitative analysis of RICM. Its use in more complex environments, with spurious reflections stemming from a variety of structures in the sample, remains however challenging. In this paper, we demonstrate two optical sectioning methods that effectively reduce such background and can be readily implemented in a conventional RICM setup: line confocal detection, and structured illumination microscopy. We characterise experimentally the benefits to image quality and demonstrate the use of the methods for quantitative imaging of complex biological and biomimetic samples: cellular membranes, thin organic films, surface biofunctionalization. We then discuss the benefits of each method and provide guidelines to arbitrate between sectioning and signal-to-noise ratio. Finally, we provide a detailed description of our experimental setup and a home-written image acquisition and processing software that should allow the interested reader to duplicate such a setup on a home-built or commercial microscope.

## Introduction

Reflection interference contrast microscopy (RICM), also known as interference reflection microscopy (IRM), has been used since the 1960s as a powerful method to image and quantify the interaction of cells with surfaces.^1,2^ It has permitted numerous studies in soft matter and biophysics, ranging from wetting to membrane dynamics and cell adhesion. It has also found numerous applications for the study of thin films,^3,4^ topology of objects near an interface^5,6^ and nano objects, a field in which the acronym iScat (“interference Scattering”) is used when optimizing for the efficient detection of light scattered by nanometric objects^7–9^ or using very high-NA objectives to maximize optical resolution.^10,11^ In short, RICM relies on the detection of the light reflected (or scattered for nanoobjects) by two or more interfaces (between regions of different optical properties) located within the coherence volume that create an interference pattern encoding information about the optical properties of, and relative distances between the interfaces.

RICM is thus a powerful technique to study thin films and object-substrate interactions, but as a wide field technique, it is sensitive to spurious reflexions outside of the volume of interest that may degrade the contrast of images and limit the sensitivity to small or weakly reflecting objects. While such background may be filtered out, physically through a phase/amplitude filter in the Fourier plane in the imaging path, or numerically at the processing stage, such approaches cannot be applied to a spatially inhomogeneous background such as created by a cell in the vicinity of the focal plane, or even the nucleus and organelles of a cell whose lower membrane is being imaged. Ideally, one would wish to detect signal stemming only from the vicinity of the coverslip - where interferences are located - similarly to the effect achieved in confocal microscopy compared to widefield fluorescence microscopy while maintaining the benefits of widefield imaging (simplicity of the optical setup, fast imaging rate).

To this aim, we demonstrate the application to RICM of two optical sectioning techniques, structured illumination microscopy (SIM)^12^ and line confocal (LC)^13^ microscopy. Optical sectioning permits to specifically detect the signal created around the focal plane, removing incoherent background to RICM images arising from out-of-focus structures. While these sectioning techniques have already been demonstrated and modelled at length for fluorescence imaging^14–18^ or to a lesser extend in reflection,^12,19^ their application to RICM has to be adapted to accommodate for partially coherent imaging and peculiar numerical aperture conditions: In RICM, it is indeed critical to tune the coherence volume in accordance with the process under study, so that interferences occur over a limited axial depth around the focal plane where the structures of interest are located. This is achieved by a mix of spatial (numerical aperture modulation) and temporal (illumination spectral width control) coherence tuning.^2^ This is in contrast with most uses of optical sectioning techniques, for which the best possible spatial resolution is sought. In particular, controlled (and generally low) illumination numerical aperture are used to facilitate the accurate retrieval of distances from the recorded interference patterns.^5,20^ Here, we therefore establish the use of sectioning techniques such as LC and SIM in the context of RICM at low illumination numerical aperture, and we detail their practical hardware and software implementations. We then present the principle of such combined techniques, and demonstrate the benefits of LC- or SI-RICM for soft matter and biophysics on three application examples: cell imaging in a crowded environment, topology of complex thin films, and monitoring of surface biofunctionalization. Finally, we discuss the benefits of optical sectioning for RICM in relation with existing alternatives.

## Materials and methods

### Optical setup

Our optical setup, apart from the digital micromirror device (DMD), is described in details elsewhere.^50^ Briefly, it is based on a modified IX71 Olympus inverted widefield microscope, with the optical turret replaced with a home-built system separating illuminating from reflected light based on polarisation. It consists of a polarisation beamsplitter cube (Thorlabs) and a custom-made optically flat achromatic quarter waveplate (Fichou, France). Illumination is provided by an incoherent plasma light source (HPLS345, Thorlabs) that is spectrally and spatially filtered using dichroic filters (FF01-457/530/628-25, or FF01-530/43-25, Semrock) and graduated diaphragms (SM1D12C, Thorlabs). Before filtering, illumination light is reflected off a high-speed DMD (V-9601, Vialux, Germany) conjugated with the image plane of the microscope objective (depending on the experiment and as mentioned in the text: UPLSAPO60XO, UPLSAPO20XO, and UPLSAPO20X Olympus, Japan). Detection is achieved on one (single colour imaging) or two (3-colour imaging) sCMOS cameras (ORCA-Flash4.0 V2 and V3, Hamamatsu, Japan). For efficient transfer of images to the acquisition computer, the two cameras are connected using dedicated boards (CameraLink pack for Flash4, Hamamatsu). For LC imaging the DMD and the cameras are synchronized using a pulse generator (4052 waveform generator, BK precision). For green and blue imaging, a home-built beamsplitting system is used to project the two images on two halves of one of the camera sensors.^50^ Each colour image is 2048×1024 pixels. The magnification between the DMD and camera planes is chosen such that one DMD pixel (10.8 *μ*m) is projected onto two camera pixels (2 × 6.5 = 13 *μ*m, corresponding to M=1.2). Details about the hardware synchronization, image acquisition program and optical alignment are given in Suppl.Fig. 1-3.

### Sample preparation

All coverslips (no 1.5, VWR) were cleaned in piranha solution and stored in ultrapure water before use.

Red blood cells were obtained from a healthy donor through the French Blood Bank, diluted in phosphate buffer saline (PBS) and let to sediment on a clean coverslip. Images were acquired within a few minutes after sample preparation.

cellulose nanocrystal/xyloglucan (CNC/XG) multilayer samples were prepared using layer-by-layer deposition of in-house-prepared CNCs from cotton linters and of XG purchased from Dainnipon (Osaka, Japan). Polyethyleneimine (Mw ≈25 000) was purchased from Sigma. A full description of the sample preparation and characterization can be found in.^37^

Supported lipi bilayer (SLB) - traptavidin (TAv) platform preparation is described in detail in.^54^ Briefly, pre-cleaned coverslips were dried, plasma treated for 3 min in water vapour and glued to a home-made teflon holder using two-component silicon glue (Twinsil-speed, Picodent, Germany). After 10 min curing, small unilamellar vesicles (SUVs, 100 *μ*g/mL) in HEPES buffer (HEPES, pH 7.4, 150 mM NaCl in ultrapure water, all from Sigma) were incubated on the coverslip for 30 min and then thoroughly rinsed with HEPES buffer. SUVs were prepared using tip sonication of a HEPES solution of 5 mol % DOPE-cap-B and 95 % DOPC (both Avanti Polar Lipids, USA). At the start of the RICM image acquisition, TAv was added at 40 *μ*g/mL. TAv was expressed and purified from *E. coli* as described previously^38^ and kindly provided by Mark Howarth (University of Cambridge. UK). For subsequent fluorescence staining, TAv was thoroughly rinsed with HEPES, and biotinylated-FITC (10 *μ*g/mL, Thermo Scientific) was incubated for 30 minutes before thorough rinsing with HEPES.

### Image acquisition

In SI-RICM, acquisition time was set between 3 and 10ms to avoid saturation of the cameras, and three successive images were taken at 10ms time difference, resulting in a total acquisition time of 30ms. In case of real-time image processing, the frame rate was limited to 20Hz (for a single camera) or 10Hz (for two cameras). For LC-RICM, the acquisition time was set by the line rate (25 *μ*s per line) and the number of simultaneous illuminated lines, ranging from 6 to 20 depending on the illumination line width. As a result, the acquisition time for each pixel was between 150 and 500 *μ*s. For comparison of fluorescence labelling quantification with RICM measurements of TAv density (Application example 3), fluorescence confocal microscopy was performed on a TCS SP8 (Leica, Germany) with a 40×, 1.30NA oil objective and a built-in autofocus. Fluorescence was excited at 488 nm with a power on the sample in the range of ≈ 2*μ*W and detected in the wavelength range of 491-629 nm with a pixel dwell time of 1.2 *μ*s and a sampling of 0.284 *μ*m/pixel. Since labelled TAv crystals produce an anisotropic signal as a function of polarization, we rotated the sample in the imaging plane and acquired 7 images over a range of 180°, and considered the maximum intensity value for each crystal.

### Image analysis

Real-time SIM images reconstruction was performed using a home-written software using LabView 11.0 (National Instrument) described in the Supplementary Materials. All other image analysis was performed using the ImageJ bundle FIJI, and reflected intensity numerical calculations were performed using Mathematica.

Reconstructed SIM images often present a small residual modulation at the pattern spatial frequency or a higher harmonic, due to slight deviations from a perfect sine pattern or to illumination fluctuations. This residual modulation is filtered out by FFT filtering centered on the pattern spatial frequency and its harmonics.

For monitoring of TAv crystal formation, prior to TAv incubation, three reference images with shifted illumination patterns were acquired and sectioned and widefield reference images were computed. 3 RICM images with shifted patterned illumination were then acquired every minute after adding TAv to the solution, and the sectioned and widefield images were computed and normalized by their reference counterparts.

For 3-colour analysis of multilayer film height (Application example 2), the contrast of each of the RICM images was adjusted between its minimum and maximum value before the three images were merged into a RGB pseudo-colour image. The corresponding colour scale was built using the formula:

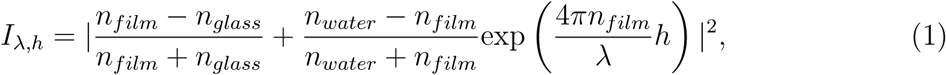

where *λ* is the wavelength, *h* is the thickness of the film, and *n_film_*, *n_glass_*and *n_water_* are the refractive indices of the multilayer film (assumed to be homogenous), coverslip and water, respectively. The curves for the 3 wavelengths (453, 532 and 632 nm) were then scaled between 0 and 1 and used to create the colour scale shown on Fig.4c. The refractive index of the film depends on its hydration, which is not a priori known. In Supplementary Fig. S4a, we show the scales obtained for a protein content of 50% (n=1.4) and 100% (n=1.47). The weak change in the colour scale illustrates the limited sensitivity to the uncertainty in the exact hydration level. In the analysis, we have then used n=1.4 (50% water content), in agreement with previous estimates of the hydration of these layers when immersed in water.^37^ Using this colour scale as a reference, the FIJI plugin Weka Trainable Segmentation^36^ is then used to classify the image as the function of pixel hue: classes of heights with 20-30 nm widths are used, and training is achieved by manually referencing pixels to each of the classes based on its hue. Manual inspection found a highest value in the image in the range 460-480nm, while the lowest values were in the range 70-80nm, setting the number of classes used for the full segmentation (16). The output of the Weka classifier is then a series of 16 probability maps for the height of each pixel (see Supplementary Fig. S5), which were combined to estimate the height through a weighted average of the 3 classes nearest to the one with the highest probability.

To quantify the roughness of the CNC/XG multilayer film, we used the widely-used arithmetic average roughness 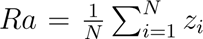, where summation is over all the N pixels of the 2D image, and *z_i_* is the estimated local thickness.

For estimating the surface density of TAv (Application example 3), the ratio between the intensities of adjacent regions within and outside of the crystalline region is computed at different positions on the sample. To relate this ratio to the local density of TAv, we use a model for the sample that includes a thin layer of medium (n=1.335, thickness 1 nm)^55,56^ above the coverslip (n=1.5196), below a lipid bilayer (n=1.4554, thickness 4.9nm),^56^ onto which a TAv layer (thickness 5 nm, in agreement with measured values for SAv^57^) has a density varying between 0 and 100% of the crystalline density (estimated to 268 ng/cm^2^ ^45^). We assume that this corresponds to a linear variation of the refractive index between 1.334 and 1.455.^58^ Coherent addition of the reflections from these different interfaces yields a TAv-concentration-dependant signal which is well described by a parabola in the range 50-100% of the crystal density (Supplementary Fig. S5). Inverting this function permits converting the experimentally measured ratio of non-crystalline-to-crystalline intensities in ratios of TAv surface densities.

## Results and discussion

### Principle and practical implementation

The proposed approaches are illustrated on Fig.1: compared to a wide-field RICM, a digital micromirror device (DMD) is inserted in the optical path and is optically conjugated with the focal plane. By applying a black-and-white amplitude mask on the device, the amplitude of the incoming light is modulated at will at frequencies up to 20 kHz and a spatial sampling in the focal plane of 216 nm (for a 60× objective, as set by the camera pixel size and DMD-to-camera magnification factor; or 648 nm with the same setup and a 20x objective). The resulting pattern is imaged on a camera and provides optical sectioning through a matching detection pattern or through post processing of the acquired image: in LC imaging (Fig.1, left), a line of controlled width is scanned across the field of view and the signal on the camera is acquired using a rolling shutter (i.e., signal acquisition starts one line after the other, instead of simultaneously for the whole camera) within a matching moving line of fixed width.^21,22^ In this case, similarly to laser scanning confocal microscopy (LSCM), optical sectioning is obtained through rejection of the defocussed reflections of the illumination line that fall mostly out of the detection line for a sufficiently large defocus.^14,18^

**Figure 1:**
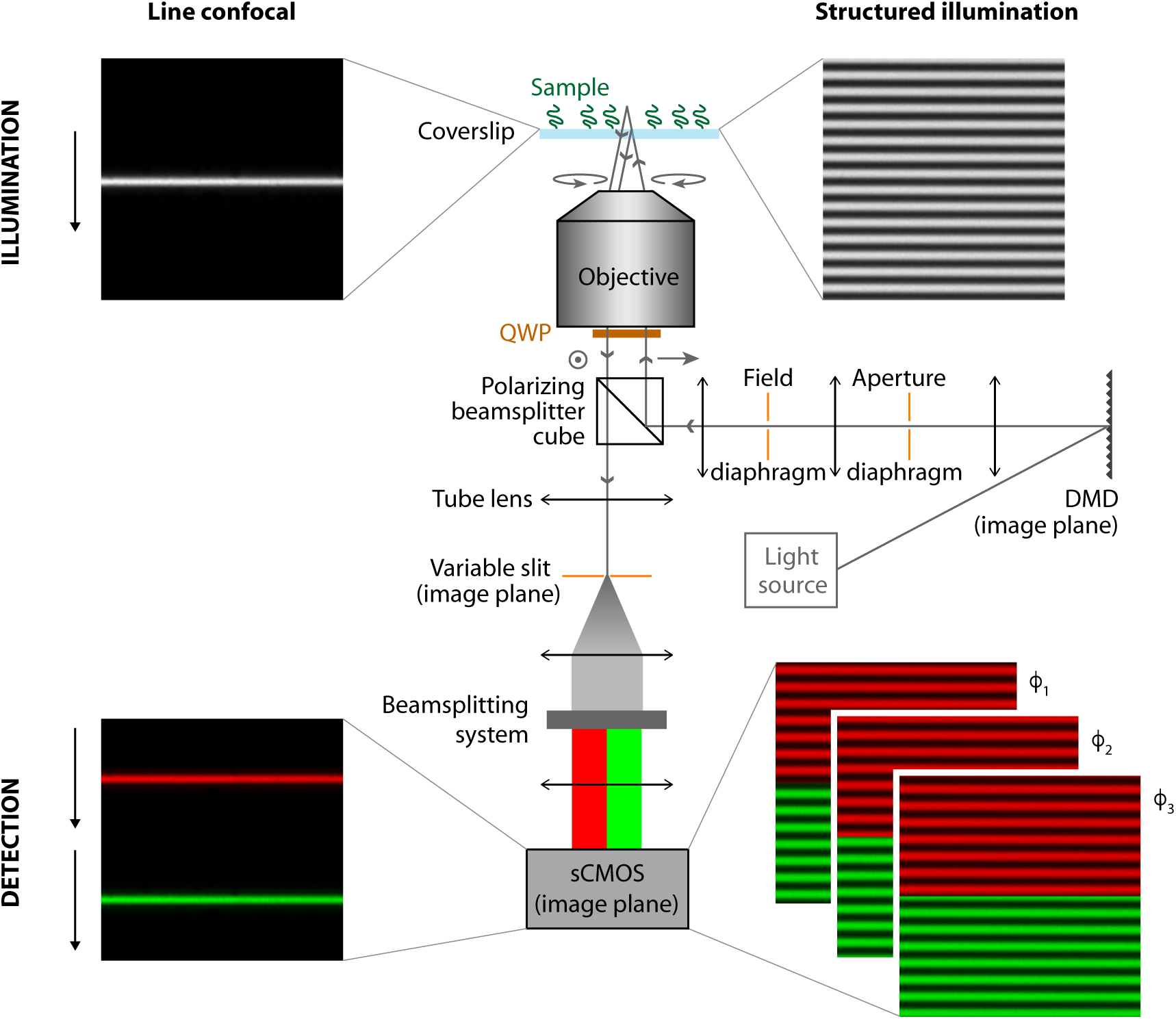
RICM with optical sectioning. Center, a schematic view of a RICM is presented, along with the illumination and detection schemes used for line confocal (LC, left) and structured illumination (SI, right) sectioning. The illumination path consists of an incoherent white light source illuminating a digital micromirror device (DMD). Its illumination cone must be large enough to fully neglect the subsequent diffraction and related chromaticism induced by the DMD. The DMD then is imaged onto the sample plane of the objective after spatial filtering through an aperture diaphragm (setting the illumination numerical aperture (INA) and hence the coherence volume within which interferences are observed^2^) and a field diaphragm (limiting the field of view). The field diaphragm is not strictly necessary as a virtual diaphragm can be set by the DMD by limiting the patterned area, but is useful for DMD alignment. The illumination beam is further filtered in polarisation using a polarising beamsplitter cube and a quarter wave plate (QWP) to separate illuminating from reflected light. The gray arrows show the polarisation of the beam at various positions in the beam path. Incoming and refected beams are drawn with an offset from the optical axis for the sake of clarity only, but are in reality incoming at the center of the objective back aperture. The reflected light is then imaged onto a sCMOS camera, either directly (not shown) or after passing through a slit and a beam splitting system to obtain simultaneously two RICM images at different wavelengths (the case of two colours is shown here as an example). Multicamera imaging is also possible to increase the number of simultaneously recorded images.

**Figure 2:**
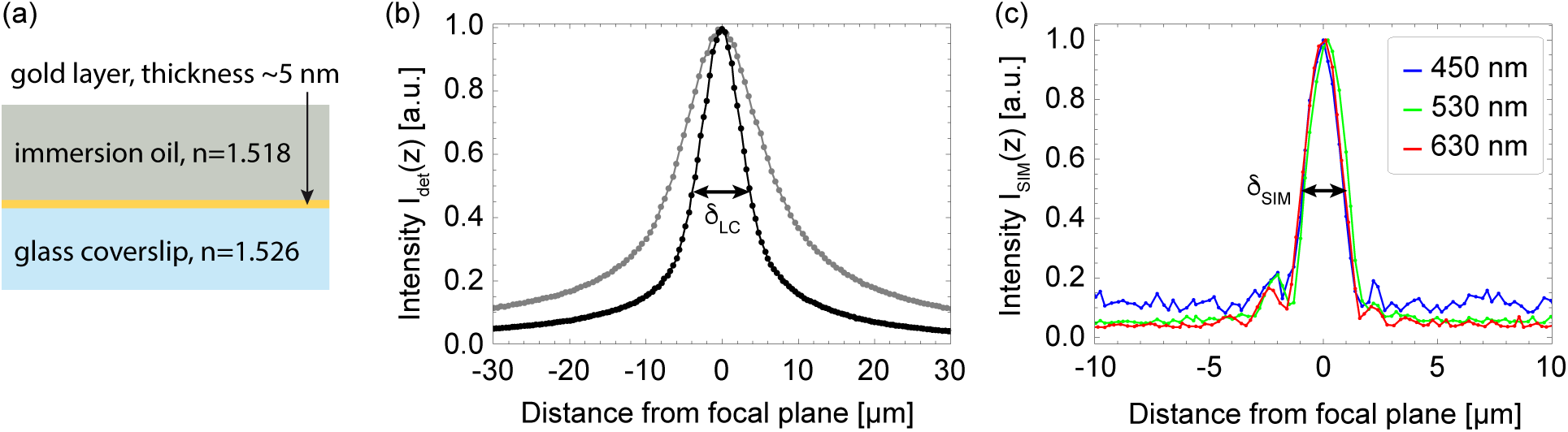
Optical sectioning in RICM. (a) Test sample used to measure the axial response of the microscope. The gold layer acts as a partially reflective layer between two semi-inifinite regions with similar refractive indices. (b) Axial response in line confocal RICM, obtained with a 20×, 0.85NA oil immersion objective and a 0.46 illumination NA (INA). For a projected line width *p* ≈ 1 *μ*m (black), *δ_LC_* ≈ 7.5 *μ*m. For a projected line width *p* ≈ 2 *μ*m (gray), *δ_LC_* ≈ 14 *μ*m. (c) Axial response in SI-RICM at three different visible wavelengths, illustrating the possibility to acquire multicolour sectioned RICM images. Profiles were acquired with a 20×, 0.75NA air objective and INA=0.46, using a pattern with period *p* ≈ 1 *μ*m. *δ_SIM_* ≈ 2 *μ*m.

Alternatively in SIM, a series of illumination structures such as grids with varying orientations and phases are projected onto the focal plane (Fig.1, right),^23,24^ and the signal is reconstructed by calculating the standard deviation of the resulting image stack.^12^ More complex algorithms can be used to improve the signal-to-noise ratio and the resolution of the reconstructions, but they all rely on the combination of at least two^25^ and up to several tens of images,^26^ thereby reducing by the same factor the frame rate of the sectioned image acquisition.^27^ The most common implementation, used in this paper, uses 3 images of a 1D sinusoidal pattern shifted between each image by one third of a period (*I*_1_, *I*_2_ and *I*_3_). While more efficient algorithms may be used to minimise the image noise and improve its resolution,^28^ we have restricted ourselves here to the canonical standard deviation calculation approach that has the benefit of simplicity and permits real time video-rate image reconstruction on a recent computer:

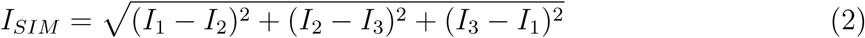

Using the same three images, a widefield image can also be reconstructed using the formula:

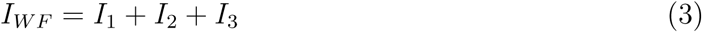

LC and SIM are complementary and can be implemented using the same experimental setup, and we detail their relative advantages and limitations in the Discussion section. Complete experimental details on their practical implementation can be found in the Methods section and Supplementary Fig. S1-2, and should allow the reader to set up an equivalent sectioning module on their microscope. Briefly, hardware TTL signals are used to synchronize the display of preloaded black and white patterns on the DMD, the acquisition of images on one or several sCMOS cameras, and if needed the motorised displacement of the sample (to acquire several fields of view or adjusting the focus, for example). For SIM, a full image is acquired on the camera with simultaneous exposure of all pixels, while LC uses progressive scan on the camera at a speed synchronised with that of the illumination line movement on the DMD. Conversely, LC images are directly displayed and recorded, while SIM images require further postprocessing to cancel the illumination patterns and obtain a sectioned image (e.g. equ. 2). A home-written LabView software is described in Supplementary Fig. S3 that permits real-time visualisation of SIM sectioned images.

To assess the sectioning ability achieved with either of these methods, we have recorded the signal reflected by a test sample composed of a glass coverslip onto which a nanometric layer of gold has been deposited that provides the optical contrast, and which is covered with immersion oil of refractive index matching that of the glass. This permits neglecting all aberrations introduced upon changes in refractive index at the interface. Here, it should be noted that we have chosen to focus on the use of small illumination numerical apertures (INA) for which illumination can be considered quasi-parallel, a widely-used approach that facilitates quantitative image analysis in RICM. INA is defined as *n* sin(*α_ill_*), with *n* the refractive index of the sample and *α_ill_*the maximum angle of the incoming illumination. A low INA also permits optimising the visibility of interferences within the focal volume, and allows visualizing objects up to ≈ 10 *μ*m above the reference surface (for INA≈ 0.25, together with the appropriate tuning of the temporal coherence).^20^ For other implementations that make use of high or even variable INAs,^5^ the formulae below still provide semi-quantitative values of pattern width and axial sectioning but deviations are expected due to high-angle propagation effects.

In low-INA implementations, the width of the patterns *p* that can be projected onto the focal volume for optical sectioning is larger than the diffraction limit of the imaging objective (typically 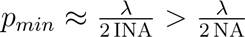, with NA the full numerical aperture of the objective). This, in turn, determines the optical sectioning thickness that is given by 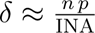. Note that for a given value of *p*, axial sectioning does not depend on the wavelength, facilitating multicolour SI-RICM. Combining both equations yields the minimal sectioning thickness 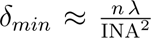 which is also the expression of the axial extension of the spatial coherence volume.^29^ A full description of the image formation process in optically sectioned RICM is beyond the scope of this paper and is the topic of another manuscript in preparation by the authors. In the case of low-INA RICM (INA in the range 0.2-0.5), one gets a shortest axial sectioning in the range 2-10 *μ*m, but values below 1 *μ*m can be achieved with higher INAs (see discussion).

While this is far from the sectioning obtained in total internal reflection fluorescence (TIRF) microscopy (a widely-used option for imaging samples close to the coverslip surface), the proposed approaches are more easily compatible with quantitative interference imaging, are easily implemented in the conventional microscope and provide a larger field of view (typically ≈ 10× larger when using a 20× objective, at the expense of a lateral resolution ≈ 1.5 – 2× larger). To demonstrate their usefulness, we provide below three case studies where optical sectioning permits the quantitative analysis of complex samples.

### Application example 1 : cell imaging in a complex environment

RICM has widely been used for marker-free quantification of cell adhesion or membrane fluctuations.^30–32^ While a powerful tool, in its classical configuration its lack of sectioning ability has hampered its use in complex environments such as in the presence of several layers of cells in which quantitative interpretation of images is challenging. In such cases, incoherent background subtraction cannot remove the spurious spatially-heterogenous, time-dependent incoherent signals arising from outside of the coherence volume.

As a proof-of-principle demonstration of optical sectioning with LC, RICM images of red blood cells in PBS sedimented on a glass slide, with and without LC sectioning, are shown on Fig.3. The lower membranes of red blood cells produce well-contrasted fringes that can be used to reconstruct their height profile. As expected, the widefield image exhibits a much higher background intensity due to incoherent parasite reflections, such as stemming from the liquid/air interface at the top of the sample. However, even when subtracting the constant offset from the images, incoherent reflections from area of cells outside the focal volume are visible in the image. In some cases, a second cell resting on a sedimented one modifies the contrast of the fringes, rendering the quantitative analysis of the latter challenging (arrows on Fig.3b). This is representative of typical physiological conditions, where the density of cells within a microvessel can reach 30-40% volume fraction.^33^ Studying cell-wall interactions at such high density would thus induce massive out-of-focus heterogeneous reflections from blood cells inside the channel.

**Figure 3:**
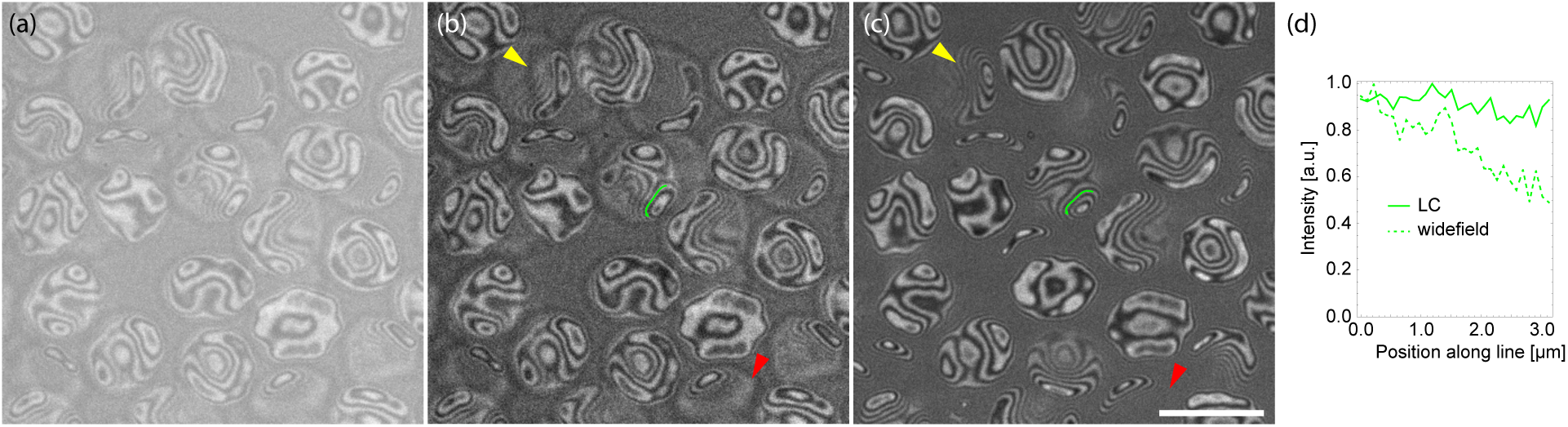
Red blood cell imaging in a crowded environment. RICM images of red blood cells sedimenting on a glass surface without (a-b) and with (c) LC sectioning. (a) Raw wide field image. (b) Same image with background removed. Outlines of cells stemming from incoherent reflections are still visible in the images (red arrowhead) and can superimpose with the interference pattern (yellow arrowhead). This results in a variable intensity of the interference fringes, as demonstrated in (d) where the profile along the green line is plotted with and without LC sectioning. Profiles were acquired with a 60×, 1.35NA air objective and INA=0.46. Scale bar, 10 *μ*m.

In contrast, the same image acquired using LC-RICM yields a much clearer image of the cell structures in the vicinity of the coverslip that provide interferential contrast. Note that the images with and without LC sectioning were acquired sequentially, so that a slight sedimentation and differences in membrane fluctuations are visible in between the two images.

### Application example 2 : quantifying the topology of complex surfaces

RICM can also be used as a non-invasive probe of the topology of surfaces decorated with organic material, including wet samples that cannot be dried without altering their structure. Compared to its derived counterpart biolayer interferometry (BLI),^34^ it can provide sub-micron resolution in the imaging plane that permits assessing not only the average layer thickness but also the thickness distribution. This is usually performed using multicolour RICM, providing unambiguous information on the layer thickness up to about 1 *μ*m.^3,4^ Quantitative analysis of the RICM signal is however often hampered by the spurious reflections from other interfaces. In particular, it is often convenient to use air objectives for intermediate characterisation steps, so that further functionalization can be performed without cleaning of the lower face of the coverslip, and without sample contamination with immersion oil. This leads to a strong reflection from the non-functionalized side of the coverslip which for samples immersed in water is typically one order of magnitude larger than the signal from the interface of interest itself. In this context, optical sectioning greatly improves the quantification of the surface topology.

As a demonstration, we have imaged a multilayer sample of alternating cellulose nanocrys-tals (CNCs) and xyloglucan (XG) particles adsorbed on a polyethileneimine (PEI) layer. XGs are the major hemicelluloses in the primary cell wall of dicotyledonous plants, while CNCs in plant cell walls combine with other biomolecules to form supramolecular assemblies reinforcing its structure. The strong interaction between CNCs and XGs permits building biomimetic nano composite films reminiscent of plant cell walls, whose thickness is controlled by the number of (CNC/XG) layers.^35^ While such layers can be characterised by AFM at small scale (typical a few *μ*m^2^), or by neutron reflectivity (with lateral averaging over typically several cm^2^),^35^ multicolour RICM with optical sectioning provides a fast, submicrometric, large field-of-view characterization of the film height that can be performed as a quality control at different stages of film deposition. We demonstrate how with a 20×, air objective, line confocal RICM permits monitoring the thickness of a PEI(CNC/XG)_5_CNC film over a 200×500 *μ*m^2^ field of view in ≈ 1s with a ≈ 500nm lateral resolution using 3 visible wavelengths (453, 532 and 632 nm) that allow reconstructing pseudo-colour images (Fig.4a). Comparison between widefield and optically sectioned images reveals a striking improvement in contrast and an increased visibility of small structures, as well as a lower sensitivity to out-of-focus impurities (Fig.4a, red arrowheads). Even after FFT filtering to remove the slowly varying background, the visibility of small structures is still hampered by noise and residual background in wide field RICM images compared to line confocal images (Fig.4b, yellow arrowheads).

**Figure 4:**
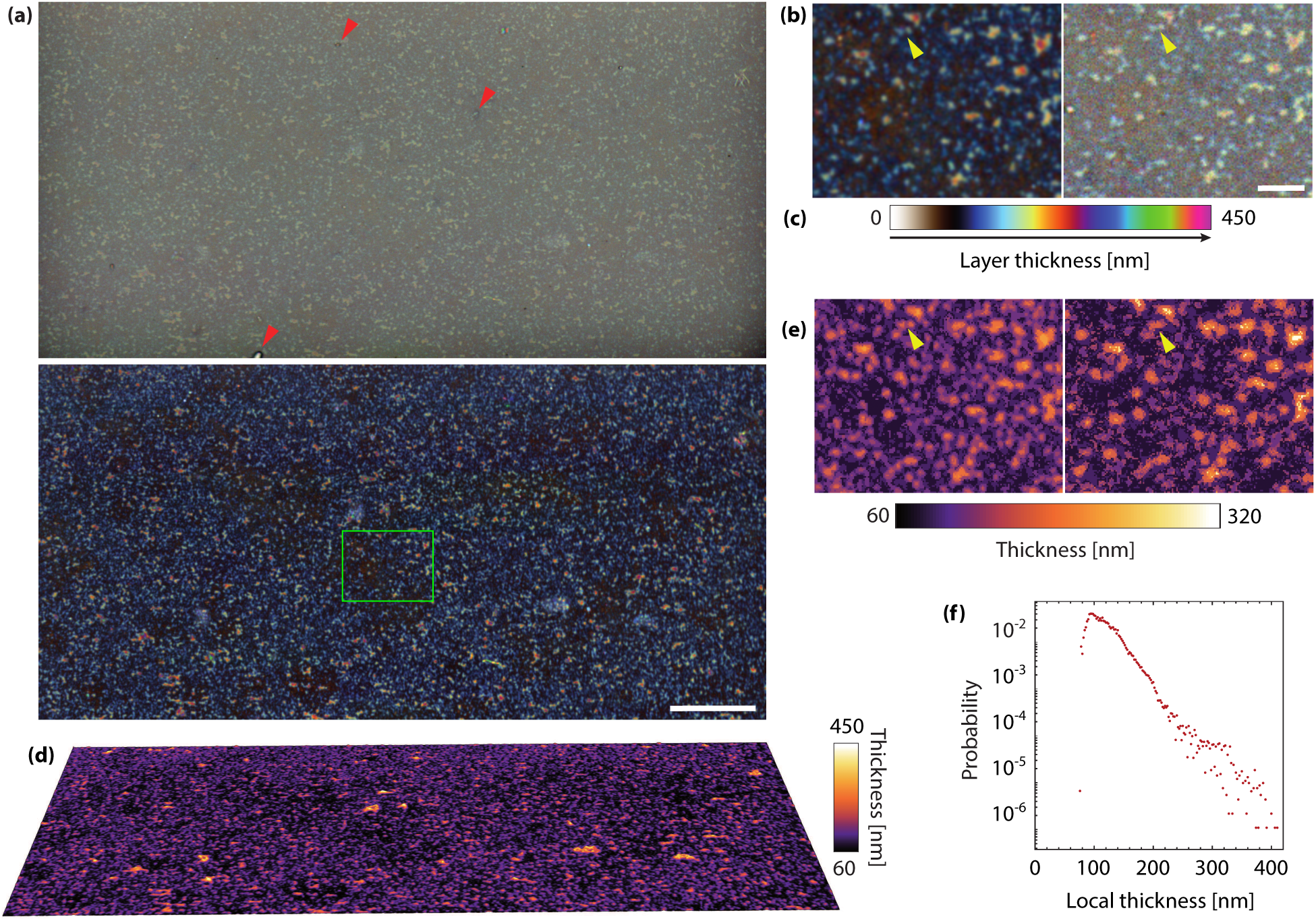
Topology of a biomimetic multilayer film. (a) A PEI(CNC/XG)_5_CNC film obtained by layer-by-layer deposition and immersed in water is visualised without (top) and with (bottom) line confocal sectioning in 3-colour RICM, yielding a false colour RGB image that informs on the topology of the film. In the absence of sectioning, a large incoherent background reduces the contrast of the signal of interest. Scale bar, 50 *μ*m. Further quantification is presented in Supplementary Fig. S6. (b) Left, zoom on the green rectangle in (a) and right, zoom on the same region of the widefield (WF) image, with a high-pass Fourier filtering applied. This preserves the RICM contrast only if assuming a zero low frequency component in the signal of interest (an assumption that fails in the case of samples thinner than about 100 nm). Scale bar, 10 *μ*m. (c) Computed hue corresponding to a given height assuming a layer refractive index of 1.4. (d,e) Reconstruction of the height profile based on the 3-colour RICM signal, on the full sectioned image (d) and on the green inset for both LC and WF RICM (e). (f) Histogram of heights measured on each pixel from the reconstruction in (d).

Furthermore, sectioned images can be used to quantitatively reconstruct the topology of the film using simple assumptions for its optical properties: assuming a homogeneous, isotropic refractive index (a reasonable assumption for a disordered film of polysaccharides) ranging from 1.47 (fully dehydrated) to 1.4 (partly hydrated, polysaccharide content 50%) we can reconstruct the variations of hue expected as a function of film thickness (Fig.4c and methods). Interestingly, the colour scale is only weakly affected by the large change in refractive index that we have considered, yielding only a ≈ 5% systematic change in the estimated thickness between these two strongly different hydration assumptions. Based on this hue scale, machine learning is used to map the 3-colour RICM image onto a series of height probability images using 20-nm steps. ^36^ These images are subsequently combined to obtain a quantitative thickness image over the whole field of view (Fig.4d and 4e, see Methods). These maps provide a wealth of information such as the average film thickness (here ≈ 121±5nm), roughness (here ≈ 24±2nm, see methods for the definition of roughness) and characteristic lateral size of height variations (here ≈ 0.65 *μ*m, close to the resolution limit ≈ 0.5 *μ*m hinting that smaller features are present but less visible or poorly resolved, in agreement with AFM images^35^), as well as a full height histogram (Fig.4f). Previous AFM characterizations have shown that the thickness of hydrated PEI(CNC/XG)_5_CNC films was 124 ± 21nm,^37^ a value with which our height measurement through confocal RICM is in excellent agreement, demonstrating the robustness of our optical measurement experimental setup and processing pipeline. As a comparison, segmentation of widefield RICM images yield noisy height maps, in particular in the 80-120 nm thickness range where the RICM contrast is low and therefore more sensitive to noise. Details of the height maps visible in the LC RICM images are therefore poorly reconstructed (Fig.4b and e, yellow arrows).

### Application example 3: probing surface biofunctionalization quantitatively

In addition to being non invasive, spatially-resolved and compatible with wet samples, RICM also permits rapid measurements which allow monitoring of sample formation or changes *in situ*. Here, we imaged the functionalization of a glass interface with a molecular platform permitting controlled presentation of biomolecules at a solid/liquid interface. The platform is composed of a supported lipid bilayer (SLB) incorporating biotinylated lipids (see Methods) that provide anchor points for tetrameric traptavidin molecules (TAv), a variant of streptavidin (SAv) with a larger binding energy.^38^ Because only two to three of the four biotin-binding sites are bound to the SLB,^39^ this platform provides a way to graft a variety of biotinylated molecules at a controlled orientation and surface density, forming well-controlled biomimetic surfaces widely used for *in vitro* studies of ligand-receptor, or interactions of cells or viruses with model cell surfaces.^40–42^ The method relies on the fact that the biotin-SAv bond is one of the strongest non-covalent bond known in nature, which is even strengthened in the synthetic variant TAv (35 *kT* /bond). However, while the SLB-SAv platform has been already widely used and characterised, and in particular the ability of mobile SAv molecules to form 2D crystals above a critical surface concentration, ^43^ its synthetic variant TAv has not been characterized so far for this type of application.

Here, we take advantage of the enhanced sensitivity of SI-RICM to monitor the adsorption of TAv to a biotinylated SLB during TAv incubation. Contrary to spectral ellipsometry, which provide a laterally-averaged quantification of the adsorbed amount of protein, SI-RICM does not straightforwardly inform on the adsorbed surface density of proteins in the regime of a fraction of a single monolayer, but provides a direct readout of the homogeneity of the functionalized surfaces. Interestingly, this sub-micron; resolution shows that in incubation conditions that provide homogeneous and mobile SAv layers, TAv forms regions of higher densities (appearing darker in RICM) surrounded by lower density areas (Fig.5). Real-time monitoring shows that this process occurs quasi-simultaneously in the whole field of view and is initiated on sub-resolution defects (green arrowheads in Fig.5a. The lateral resolution is ≈ 250 nm, using a 60×, 1.35NA oil objective) in the bilayer (see Supplementary Movie 1). By analogy with SAv, for which literature extensively documents the formation of 2D crystals on lipid layers,^43–46^ and since when labelled (see below), dense areas display an anisotropic in-plane response and an absence of mobility despite the fluidity of the SLB (confocal imaging and FRAP, not shown), we similarly attribute the dense phase to the formation of 2D crystals.

**Figure 5:**
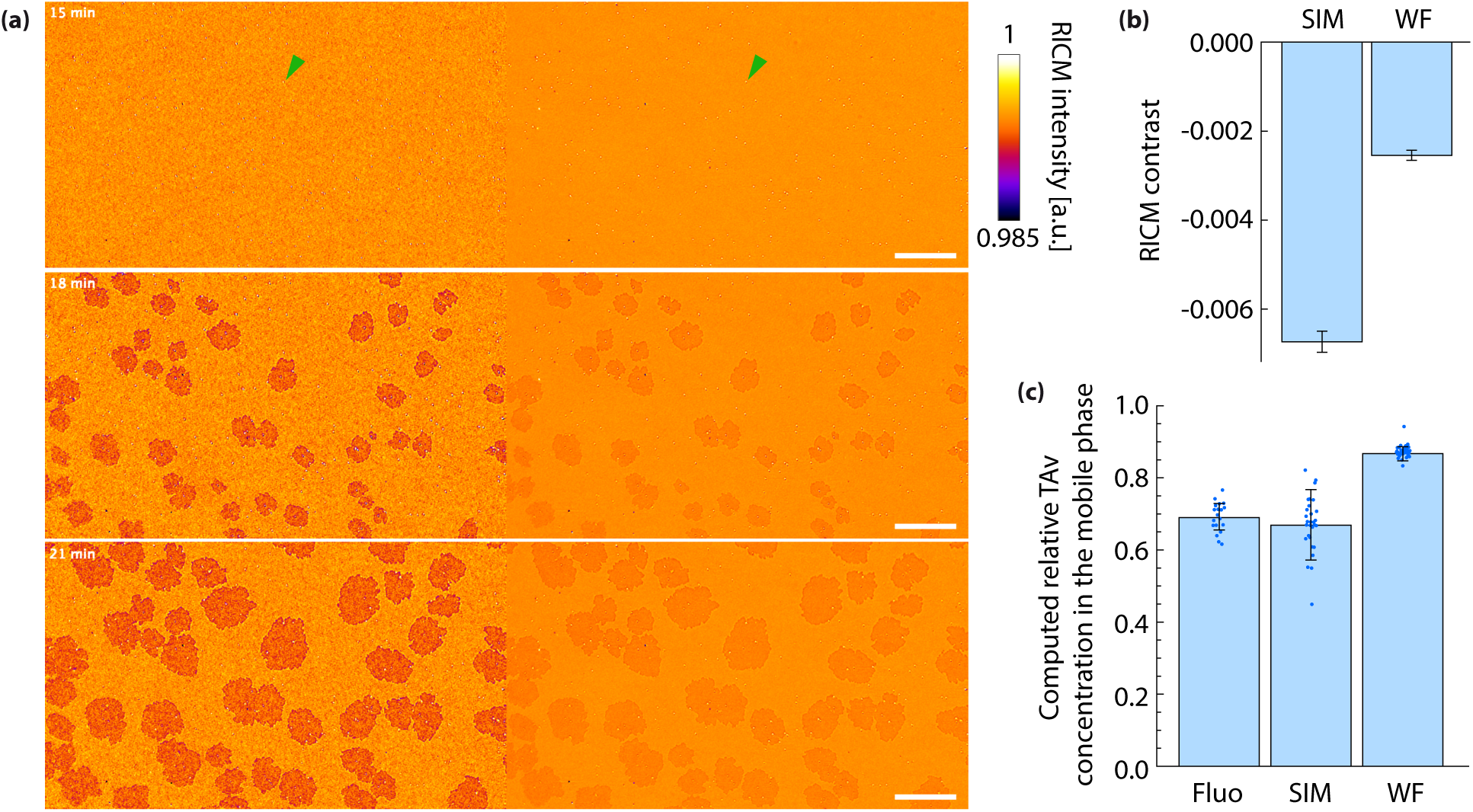
In situ, non-invasive monitoring of TAv adsorption to a biotinylated surface. (a) 532-nm SI (left) and widefield (right) RICM images of a biotinylated SLB incubated with TAv. Crystals start to form on the surface after ≈ 17 min incubation, with a contrast that remains constant over the whole incubation phase. Arrowheads point to defects in the bilayer, such as unbursts lipid vesicles, that act as nucleating centers for crystallization. Scale bars, 20 *μ*m. (b) Relative change in intensity in the crystalline phase compared to the mobile phase with (SIM) and without (WF) SI sectioning, illustrating the gain in contrast thanks to optical sectioning. Error bars are standard errors of the mean on n=24 independent measurements on different crystals of the same sample (SIM: −6.7 × 10^−3^ ± 4.3 × 10^−4^; WF: −2.35×10^−3^±6.9×10^−5^). (c) Computed concentration of TAv in the mobile phase compared to the crystalline phase using fluorescence confocal microscopy (Fluo), SI and widefield (WF) RICM. SI-RICM is in good agreement with confocal measurements, while widefield RICM underestimates the difference in surface concentration between the two phases. Error bars are standard deviations on n=16 (confocal) or n=24 (RICM) independent measurements on different crystals of the same sample, represented as blue dots. The same sample was measured in RICM and fluorescence to allow direct comparison of the measured densities.

This application highlights the extreme sensitivity of SI-RICM to minute variations of the surface reflectivity which is modulated here by only 0.6%. Importantly, optical sectioning permits analyzing the changes in reflectivity to extract quantitative information about the sample. Here, we observe first that the contrast between the high- and low-density phases is constant over time, indicating that once a sufficient surface density is reached, additional binding of TAv does not increase further the density of the mobile phase, but instead induces growth of the crystalline phase. Furthermore, modelling the optical properties of the SLB-TAv platform and assuming that the TAv crystalline phase density is identical to that of SAv (34 nm^2^/molecule, or 4.9 pmol/cm^2^, for crystallization conditions similar to ours^44,45^), we can estimate the density of TAv in the disordered phase (see Methods). We find that the mobile phase is about 30-35% less dense than the crystalline phase (corresponding to 3.3±0.1 pmol/cm^2^), while analyzing widefield images provides a relative difference of only 13% (corresponding to 4.3±0.05 pmol/cm^2^). This stems from the lesser contrast of the crystals in the presence of an unknown incoherent background.

To validate the value obtained by SI-RICM, we incubated the platform further with biotinylated-FITC and imaged the sample using fluorescence confocal microscopy: comparing the relative fluorescence intensities in the crystalline and mobile regions confirms a relative change of ≈ 30%, highlighting the benefits of optical sectioning for the quantitative analysis of RICM images. Comparing RICM with quantitative confocal fluorescence imaging demonstrates that while SIM increases the photon noise in the sectioned images (and hence decreases the precision of measurements), it removes systematic biaises in intensity quantification, thereby improving RICM accuracy. Note that while fluorescent tagging provides a more straightforward measure of TAv surface density, it can only be used after the end of incubation, and prevents further functionalization by biotinylated molecules of interest. This highlights the benefits of SI-RICM for investigating surface functionalization.

### Comparison between LC and SIM RICM, and with other sectioning approaches

In this paper, we have demonstrated two complementary methods for obtaining optical sectioning in a RICM through patterning of the illumination light. We have also illustrated the use of these methods for the quantitative monitoring of soft matter samples with high temporal and spatial resolution. Both methods improve the sensitivity of wide-field RICM at the expense of a moderate increase in the imaging system complexity, and require the use of a light patterning device, and of a light source powerful enough to afford the loss of ≈ 50% of the illumination power upon patterning (due to reflection losses on the DMD). However, while reaching optimal axial sectioning requires particular care in the design and alignment of the optical system, the small INAs often used in RICM minimize issues of chromatic aberrations and field aplanetism compared to, e.g., TIRF microscopy. This makes such a system particularly simple to install, and we estimate that a student with experience in optics can setup an equivalent system in about two weeks. To this aim, we provide detailed information about the setup (Fig.1 and Supplementary Fig. S1) and its control and synchronization (Supplementary Fig. S2), to help the interested reader implement a similar sectioning method on their microscope.

To better comprehend the situations in which LC or SIM sectioning are particularly useful, we will first detail the limitations of both methods. LC sectioning, first, requires the patterning device to operate at high speed (typically several kHz to tens of kHz, setting the line rate of image acquisition), and a sCMOS camera with a successive line detection scheme that can be synchronized with the patterning device. This requires particular alignment care when multicolour imaging is sought. In addition, in this approach, the illuminating beam is confined along a single, instead of two, spatial dimensions compared to point-scanning confocal imaging and the resulting signal decays relatively slowly with the distance to the focal plane.^14,47^ Its efficiency to cut out the signal reflected from a very bright interface close to the focal plane is thus more limited than that of SIM (see Fig.2b and 2c). Finally, the sectioning in LC is obtained at the expense of a reduction of the integration time per pixel (typically by a factor *N_ill_/N_tot_*, with *N_tot_* the total number of DMD lines imaged on the camera, and *N_ill_* the number of DMD lines simultaneously illuminating the sample), and hence of the brightness of images. As a result, the benefit of optical sectioning to remove out-of-focus noise may be offset by the increased photon noise on the resulting image, unless long integration times and hence low frame rates are used. In low-INA RICM, most light sources (e.g. high-power LED, halogen) permit exploiting the full dynamic range of the camera with exposure time in the ms range. To maintain a frame rate of 10-100 Hz with an optimal sectioning, the exposure time in LC imaging should be reduced to 20-300 *μ*s (depending on the number of line on the camera and its maximal frame rate), corresponding to a factor 10-100 compared to WF. This translates into a 3-10 fold decrease in the signal to noise ratio (SNR), unless the power of the illumination light source can be increased by a factor *N_tot_/N_ill_*. It should be noted that the decrease in integration time corresponds to an equivalent decrease in sample exposure, so that longer integration or higher intensity would not induce additional light-induced detrimental effects. An alternative is to enhance intensity through physical scanning of the image of the light source focussed onto a thin line, although this approach makes synchronization between illumination and detection scans challenging and only partially mitigates the light loss for an incoherent, high-NA light source as mostly used in widefield RICM. Another mitigation strategy, when possible is to increase the INA: in the example of Fig.3, INA can be increased up to 1.35 which, for a flat illumination profile of the back aperture of the objective, corresponds to an increase of (1.35/0.46)^2^ ≈ 9 in intensity, reducing the drop in RICM signal to a factor 1-10, and that in the SNR to a factor 1-3. Conversely, the advantage of LC, however, is that if this drop in SNR is acceptable, or if the illumination intensity can be increased, the imaging frame rate is unaffected compared to WF imaging and sectioning is obtained through a single image. As a result, fast processes are imaged without loss of temporal dynamics, and without reconstruction artefacts. In addition, if illumination intensity can be increased, LC rejection of out-of-focus background permits taking advantage of the full dynamics of the camera.

In contrast, SIM is compatible with most cameras and patterning devices provided that image acquisition can be synchronized with light patterning at the frame rate (instead of the line rate). This also permits the use of slower, less expensive patterning devices. In addition, sectioning is improved with SIM compared to LC: SIM sectioning can be compared to point-scanning confocal, with intensity dropping sharply outside of the focal volume. In contrast, LC relies on confinement in one spatial dimension only and the intensity therefore decreases more slowly as 1/*z* (with *z* the distance from the focal plane), producing a further-reaching tail in the axial intensity distribution (Fig.1). The main drawback of SIM, though, is that contrary to LC it yields raw images with a very strong modulation that prevents observing the sample in real time, unless quasi-real-time processing is performed to display a sectioned image. In Supplementary Fig. S3, we provide the interested reader with a guide on how to set up such real-time processing, which permits mitigating this limitation. Another limitation of SIM is that the nonlinear processing used to retrieve the sectioned image also results in an increase of the residual photon noise in the sectioned image, both from the coherent signal and from the incoherent background (compare e.g. the SI- and WF-RICM images in Fig.5a). While the noise is generally low for the large signals achieved in reflection imaging (in contrast to fluorescence imaging), a consequence is that removing a known background through background subtraction proves more efficient than sectioning. This is also true for a constant background that can be recorded separately, for example stemming from fixed, out-of-focus reflexions in the microscope that are imperfectly suppressed by the quarter waveplate (these spurious reflexions are experimentally negligible in our optical setup).

For both methods, it thus appear that they offer a particular benefit when subtraction of a known background is not possible, which occurs in a wide range of situations encountered practically: changing or unknown background, complex incoherent reflections from upper parts of the object under investigation, etc. Indeed, background subtraction requires that it is possible to accurately record a background image, which is most often difficult to distinguish from the homogeneous reflexion of the coverslip (and hence biases quantitative reflection variation measurements, see Application 3) or might change over time (see Application 1). When such background is large, LC and SIM are also particularly useful for achieving well-contrasted imaging of the sample in real time when exploring a sample in which the background changes with the position and cannot be efficiently subtracted: microfluidic channel, well with a curved upper free surface for the liquid, etc. This provides valuable help to adjust imaging parameters in real time (e.g. focus) and select easily relevant areas of the sample under investigation.

Table 1 summarizes the comparative benefits of each background removal method.

**Table 1:** Summary of background removal options for widefield RICM. Comparative benefits of the different methods for background removal. Frame rate refers to the case where processing is performed offline, except for line confocal. Image brightness considers identical acquisition time per image and illumination power, and can be mitigated through adjustment of the illumination power (line above). Signal to noise ratio is given for a comparable illumination power and acquisition time for the three methods, and can be reduced by increased exposure, averaging or illumination power. Movement artifacts refer to artifacts induced by the background removal operation. Additionally, distortions or blurring can occur for movements faster than the pixel exposure time.

|  | **Background subtraction** | **Line confocal** | **Structured illumination** |
| --- | --- | --- | --- |
| Known background required | Yes | No | No |
| Frame rate | max | max | max/3 |
| Illumination power | standard | $\approx \times 100$ | $\approx \times 2$ |
| Image brightness | standard | or $\approx /100$ | or $\approx /2$ |
| Background noise | $\sqrt{I_{background}}$ | 0 | $\frac{2}{\sqrt{3}}\sqrt{I_{background}}$ |
| Movement artifact | - | - | + |

Other approaches have aimed at improving the contrast of RICM images. Interferometric scattering (iScat) provides unprecedented contrast on point-like objects such as individual proteins through a combination of spatial and temporal filtering of the signal.^8^ While this method is orders of magnitude more efficient for picking out unlabelled nanometric objects down to the single macromolecule, its use to quantify extended structures requires a complex optical setup and image processing pipeline.^10^ Similarly, label-free evanescent reflection imaging (epi-EM),^48^ which uses an optical configuration similar to TIRF, relies on spatial filtering of the small-angle component of the illumination and detection light. While it permits accurate quantification of the height of relatively well-known objects above a substrate, the use of TIRF objectives limits its field of view and the reconstruction of height profile is quite sensitive to the assumptions on refractive indices in the sample. Both iScat and epi-EM can thus be seen as complementary methods to investigate fine details of relatively well-known samples, while LC- or SI-RICM provides a wide field-of-view mapping of less well-known samples that supports more straightforward quantitative analysis. Finally, it is important to underline that while we have focussed in this paper on low-INA implementation of RICM, the sectioning methods presented here are fully compatible with high-INA imaging, as has been extensively demonstrated in fluorescence microscopy:^49^ when using the full numerical aperture of the objective, the best achievable axial resolution in SIM is actually the same as in, e.g. confocal iScat.

We note for our soft matter readership, however, that high-INA diffraction-limited axial sectioning can be detrimental in some cases: it is a general rule that the better the resolution or sectioning, the greater the constraints on the defects of the optical system. A common example is the field curvature, due to which areas further away from the center of the field of view are slightly out of focus. Whereas the effect of a slight defocus on RICM signals can often be taken into account for quantitative analysis, a loss of signal due to axial sectioning reduces the accessible field of view, all the more because it occurs for a defocus twice smaller than interference losses due to the double pass effect through the optical system (illumination and detection). While for some applications, it may act as a spatial filtering ensuring that all the analyzed signals are strictly in focus, for cases where large fields of view are required an increased depth of focus may be beneficial, as demonstrated in Application 2. Another example is the case of objects interacting at varying distances of the reference surface, for which increased depth of focus is essential to maintain the visibility of the fringes over the range of heights under investigations.^20,50^ Interestingly, in LC and SIM RICM, the sectioning power can be straightforwardly adapted through the modulation of the width of the pattern *p*, similarly to the change in pinhole size in confocal microscopy. This flexibility ensures that LC or SIM should provide the user with the best compromise between out-of-focus rejection and sample visualization.

Another option to increase the contrast of interest is the combination of RICM with custom-designed coverslip coated with multiple layers of transparent material which, upon illumination at the right angle and wavelength, provide a quantitative, dark field image of material deposited at the interface.^51^ This method yields highly contrasted images without the need for further image processing of thin deposits of material at the coverslip surface, and has been demonstrated e.g. for mapping bacterial trails without staining^52^ or monitoring surface functionalization with supported lipid bilayers.^53^ It relies on the use of specific coverslips that considerably increase the cost of experiments and can also limit flexibility in sample preparation (e.g., surface biofunctionalization with thiols requiring addition of a gold layer on the coverslip), and imposes some constrains on the optical system (INA, illumination wavelength). Conversely, LC- and SI-RICM do not impose any constraint on the type of sample under investigation beyond that of classical RICM (i.e., having a reference surface providing one of the reflected light waves creating the interference pattern): inorganic (gold, silica, anti-reflective or reflection-enhancing coatings) or organic (e.g. surface biofunctionalization, thin adhesive or anti-adhesive coatings) deposits can be used without affecting the axial sectioning. In addition, the sample can be included in more complex environments such as microfluidic devices, well plates, etc. Again, both approaches can be seen as complementary, depending on the sample under investigation.

Finally, we underline here that both LC and SIM are compatible with multicolour RICM, which facilitates quantitative analysis of samples whose structure is not known precisely,^4,5^ and with fluorescence imaging, that would also benefit from the same optical sectioning. Since the methods do not impose any constraint on the illumination wavelength, any fluorophore can be used for combined RICM/fluorescence imaging. In fact, setting up the sectioning approaches proposed in this paper paves the way to the use of numerous techniques using patterned illumination, e.g. in the photoactivation or optogenetic fields. Reciprocally, already installed commercial patterning modules might permit SI-RICM without the need of additional hardware. Together, these points make the proposed approaches highly versatile for imaging soft matter or biological samples.

## Conclusion

In this paper, we have demonstrated the use of two optical sectioning methods, line confocal and structured illumination, to provide widefield RICM with axial sectioning, thereby improving the contrast of structures of interest in a crowded environment or in the presence of a strong, unknown incoherent background. Both methods are demonstrated on application examples relevant for biophysical and soft matter studies, and quantification of the optically-sectioned RICM images is demonstrated. The flexibility of these methods should make them useful in a number of experimental situations, and we provide the readers with a detailed guide on how to setup such sectioning methods to make it widely accessible to non-specialists.

## Supporting Information

Additional details on the setup (optical alignment and electronic control) and its control software, and on data analysis.

Movie of traptavidin crystallization.

## Acknowledgments

The authors thank Gwennou Coupier for help with the red blood cell experiment and Mark Howarth for the kind provision of the traptavidin. This work was supported by ANR grant HiTrac (ANR-19-CE42-0010), CNRS MITI grant TOC-SRIM and UGA IRGA grant SRIM-FAST to D.D., ANR grant MULTIMOD (ANR-19-CE37-0007) to C.V., ANR grant LEARN (ANR-22-CE16-0013) to C.V. and Y.C., LabEx Tec21 (Investissements d’Avenir, ANR-11-LABX-0030), and the BBSRC (Partnering Award BB/W018500/1; to R.P.R.).

## References

(1) Curtis, A. S. G. The Mechanism of Adhesion of Cells to Glass: A Study by Interference Reflection Microscopy. The Journal of Cell Biology 1964, 20, 199–215.

(2) Limozin, L.; Sengupta, K. Quantitative Reflection Interference Contrast Microscopy (RICM) in Soft Matter and Cell Adhesion. ChemPhysChem 2009, 10, 2752–2768.

(3) Huerre, A.; Jullien, M.-C.; Theodoly, O.; Valignat, M.-P. Absolute 3D Reconstruction of Thin Films Topography in Microfluidic Channels by Interference Reflection Microscopy. Lab on a Chip 2016, 16, 911–916.

(4) Varma, S.; Bureau, L.; Débarre, D. The Conformation of Thermoresponsive Polymer Brushes Probed by Optical Reflectivity. Langmuir 2016, 32, 3152–3163.

(5) Dejardin, M.-J.; Hemmerle, A.; Sadoun, A.; Hamon, Y.; Puech, P.-H.; Sengupta, K.; Limozin, L. Lamellipod Reconstruction by Three-Dimensional Reflection Interference Contrast Nanoscopy (3D-RICN). Nano Letters 2018, 18, 6544–6550.

(6) Kohler, F.; Pierre-Louis, O.; Dysthe, D. K. Crystal Growth in Confinement. Nature Communications 2022, 13, 6990.

(7) Jacobsen, V.; Stoller, P.; Brunner, C.; Vogel, V.; Sandoghdar, V. Interferometric Optical Detection and Tracking of Very Small Gold Nanoparticles at a Water-Glass Interface. Optics Express 2006, 14, 405.

(8) Young, G.; Kukura, P. Interferometric Scattering Microscopy. Annual Review of Physical Chemistry 2019, 70, 301–322.

(9) Talà, L.; Fineberg, A.; Kukura, P.; Persat, A. Pseudomonas Aeruginosa Orchestrates Twitching Motility by Sequential Control of Type IV Pili Movements. Nature microbiology 2019, 4, 774–780.

(10) Küppers, M.; Albrecht, D.; Kashkanova, A. D.; Lühr, J.; Sandoghdar, V. Confocal Interferometric Scattering Microscopy Reveals 3D Nanoscopic Structure and Dynamics in Live Cells. Nature Communications 2023, 14, 1962.

(11) Mazaheri, M.; Kasaian, K.; Albrecht, D.; Renger, J.; Utikal, T.; Holler, C.; Sandoghdar, V. iSCAT Microscopy and Particle Tracking with Tailored Spatial Coherence. Optica 2024, 11, 1030.

(12) Neil, M. A. A.; Juskaitis, R.; Wilson, T. Method of Obtaining Optical Sectioning by Using Structured Light in a Conventional Microscope. Optics Letters 1997, 22, 1905–1907.

(13) Koester, C. J. Scanning Mirror Microscope with Optical Sectioning Characteristics: Applications in Ophthalmology. Applied Optics 1980, 19, 1749.

(14) Wolleschensky, R.; Zimmermann, B.; Kempe, M. High-Speed Confocal Fluorescence Imaging with a Novel Line Scanning Microscope. Journal Of Biomedical Optics 2006, 11, 064011–14.

(15) Dusch, E.; Dorval, T.; Vincent, N.; Wachsmuth, M.; Genovesio, A. Three-dimensional Point Spread Function Model for Line-scanning Confocal Microscope with High-aperture Objective. Journal of Microscopy 2007, 228, 132–138.

(16) Karadaglíc, D.; Wilson, T. Image Formation in Structured Illumination Wide-Field Fluorescence Microscopy. Micron 2008, 39, 808–818.

(17) Wilson, T. Optical Sectioning in Fluorescence Microscopy. Journal of Microscopy 2011, 242, 111–116.

(18) Mertz, J. Introduction to Optical Microscopy. 2019.

(19) Karadaglíc, D. Image Formation in Conventional Brightfield Reflection Microscopes with Optical Sectioning Property via Structured Illumination. Micron 2008, 39, 302–310.

(20) Schilling, J.; Sengupta, K.; Goennenwein, S.; Bausch, A. R.; Sackmann, E. Absolute Interfacial Distance Measurements by Dual-Wavelength Reflection Interference Contrast Microscopy. Physical Review E 2004, 69, 021901.

(21) Mei, E.; Fomitchov, P.; Graves, R.; Campion, M. A Line Scanning Confocal Fluorescent Microscope Using a CMOS Rolling Shutter as an Adjustable Aperture. Journal of Microscopy 2012, 247, 269–276.

(22) Muller, M. S.; Elsner, A. E.; Ozawa, G. Y. Non-Mydriatic Confocal Retinal Imaging Using a Digital Light Projector. Ophthalmic Technologies XXIII. 2013; p 85670Y.

(23) Dan, D.; Lei, M.; Yao, B.; Wang, W.; Winterhalder, M.; Zumbusch, A.; Qi, Y.; Xia, L.; Yan, S.; Yang, Y.; Gao, P.; Ye, T.; Zhao, W. DMD-based LED-illumination Super-resolution and Optical Sectioning Microscopy. Scientific Reports 2013, 3, 1116.

(24) Saxena, M.; Eluru, G.; Gorthi, S. S. Structured Illumination Microscopy. Advances in Optics and Photonics 2015, 7, 241–275.

(25) Lim, D.; Chu, K. K.; Mertz, J. Wide-Field Fluorescence Sectioning with Hybrid Speckle and Uniform-Illumination Microscopy. Optics Letters 2008, 33, 1819–1821.

(26) Ventalon, C.; Heintzmann, R.; Mertz, J. Dynamic Speckle Illumination Microscopy with Wavelet Prefiltering. Optics Letters 2007, 32, 1417–1419.

(27) Mertz, J. Optical Sectioning Microscopy with Planar or Structured Illumination. Nature Methods 2011, 8, 811–819.

(28) Gustafsson, M. G. L. Surpassing the Lateral Resolution Limit by a Factor of Two Using Structured Illumination Microscopy. Journal of Microscopy 2000, 198, 82–87.

(29) Rädler, J.; Sackmann, E. Imaging Optical Thicknesses and Separation Distances of Phospholipid Vesicles at Solid Surfaces. J. Phys. II France 1993, 3, 727–748.

(30) Hategan, A.; Sengupta, K.; Kahn, S.; Sackmann, E.; Discher, D. E. Topographical Pattern Dynamics in Passive Adhesion of Cell Membranes. Biophysj 2004, 87, 3547–3560.

(31) Huang, Z.-H.; Massiera, G.; Limozin, L.; Boullanger, P.; Valignat, M.-P.; Viallat, A. Sensitive Detection of Ultra-Weak Adhesion States of Vesicles by Interferometric Microscopy. Soft Matter 2010, 6, 1948.

(32) Norman, L. L.; Brugés, J.; Sengupta, K.; Sens, P.; Aranda-Espinoza, H. Cell Blebbing and Membrane Area Homeostasis in Spreading and Retracting Cells. Biophysical Journal 2010, 99, 1726–1733.

(33) Baskurt, O. K.; Hardeman, M. R.; Rampling, M. W. Handbook of Hemorheology and Hemodynamics; IOS press, 2007; Vol. 69.

(34) Concepcion, J. et al. Label-Free Detection of Biomolecular Interactions Using Bio-Layer Interferometry for Kinetic Characterization. Combinatorial Chemistry & High Throughput Screening 2009, 12, 791–800.

(35) Jean, B.; Heux, L.; Dubreuil, F.; Chambat, G.; Cousin, F. Non-Electrostatic Building of Biomimetic Cellulose-Xyloglucan Multilayers. Langmuir 2009, 25, 3920–3923.

(36) Arganda-Carreras, I.; Kaynig, V.; Rueden, C.; Eliceiri, K. W.; Schindelin, J.; Cardona, A.; Sebastian Seung, H. Trainable Weka Segmentation: A Machine Learning Tool for Microscopy Pixel Classification. Bioinformatics 2017, 33, 2424–2426.

(37) Navon, Y. Interaction of Plant Cell Wall Building Blocks: Towards a Bioinspired Model System. PhD thesis, Université Grenoble Alpes; Ben-Gurion university of the Negev, Grenoble, France, 2020.

(38) Chivers, C. E.; Crozat, E.; Chu, C.; Moy, V. T.; Sherratt, D. J.; Howarth, M. A Streptavidin Variant with Slower Biotin Dissociation and Increased Mechanostability. Nature Methods 2010, 7, 391–393.

(39) Dubacheva, G. V.; Araya-Callis, C.; Geert Volbeda, A.; Fairhead, M.; Codée, J.; Howarth, M.; Richter, R. P. Controlling Multivalent Binding through Surface Chemistry: Model Study on Streptavidin. Journal of the American Chemical Society 2017, 139, 4157–4167.

(40) Wu, J.-C.; Tseng, P.-Y.; Tsai, W.-S.; Liao, M.-Y.; Lu, S.-H.; Frank, C. W.; Chen, J.-S.; Wu, H.-C.; Chang, Y.-C. Antibody Conjugated Supported Lipid Bilayer for Capturing and Purification of Viable Tumor Cells in Blood for Subsequent Cell Culture. Biomaterials 2013, 34, 5191–5199.

(41) Migliorini, E.; Thakar, D.; Sadir, R.; Pleiner, T.; Baleux, F.; Lortat-Jacob, H.; Coche-Guerente, L.; Richter, R. P. Well-Defined Biomimetic Surfaces to Characterize Glycosaminoglycan-Mediated Interactions on the Molecular, Supramolecular and Cellular Levels. Biomaterials 2014, 35, 8903–8915.

(42) Di Iorio, D.; Verheijden, M. L.; Van Der Vries, E.; Jonkheijm, P.; Huskens, J. Weak Multivalent Binding of Influenza Hemagglutinin Nanoparticles at a Sialoglycan-Functionalized Supported Lipid Bilayer. ACS Nano 2019, 13, 3413–3423.

(43) Calvert, T. L.; Leckband, D. Two-Dimensional Protein Crystallization at Solid-Liquid Interfaces. Langmuir 1997, 13, 6737–6745.

(44) Wang, S.-W.; Robertson, C. R.; Gast, A. P. Two-Dimensional Crystallization of Streptavidin Mutants. The Journal of Physical Chemistry B 1999, 103, 7751–7761.

(45) Yamamoto, D.; Nagura, N.; Omote, S.; Taniguchi, M.; Ando, T. Streptavidin 2D Crystal Substrates for Visualizing Biomolecular Processes by Atomic Force Microscopy. Biophysical Journal 2009, 97, 2358–2367.

(46) Fukuto, M.; Wang, S.; Lohr, M. A.; Kewalramani, S.; Yang, L. Effects of Surface Ligand Density on Lipid-Monolayer-Mediated 2D Assembly of Proteins. Soft Matter 2010, 6, 1513–1519.

(47) Tal, E.; Oron, D.; Silberberg, Y. Improved Depth Resolution in Video-Rate Line-Scanning Multiphoton Microscopy Using Temporal Focusing. Optics Letters 2005, 30, 1686–1688.

(48) Bon, P.; Barroca, T.; Lévèque-Fort, S.; Fort, E. Label-free Evanescent Microscopy for Membrane Nano-tomography in Living Cells. Journal of Biophotonics 2014, 7, 857–862.

(49) Gustafsson, M. G. L.; Shao, L.; Carlton, P. M.; Wang, C. J. R.; Golubovskaya, I. N.; Cande, W. Z.; Agard, D. A.; Sedat, J. W. Three-Dimensional Resolution Doubling in Wide-Field Fluorescence Microscopy by Structured Illumination. Biophysj 2008, 94, 4957–4970.

(50) Davies, H. S.; Baranova, N. S.; El Amri, N.; Coche-Guérente, L.; Verdier, C.; Bureau, L.; Richter, R. P.; Débarre, D. An Integrated Assay to Probe Endothelial Glycocalyx-Blood Cell Interactions under Flow in Mechanically and Biochemically Well-Defined Environments. Matrix biology : journal of the International Society for Matrix Biology 2019, 78–79, 47–59.

(51) Ausserré, D.; Valignat, M. P. Surface Enhanced Ellipsometric Contrast (SEEC) Basic Theory and *λ*/4 Multilayered Solutions. Optics Express 2007, 15, 8329–8339.

(52) Ducret, A.; Valignat, M.-P.; Mouhamar, F.; Mignot, T.; Theodoly, O. Wet-Surface– Enhanced Ellipsometric Contrast Microscopy Identifies Slime as a Major Adhesion Factor during Bacterial Surface Motility. Proceedings Of The National Academy Of Sciences Of The United States Of America 2012, 109, 10036–10041.

(53) Gunnarsson, A.; Bally, M.; Jönsson, P.; Médard, N.; Höök, F. Time-Resolved Surface-Enhanced Ellipsometric Contrast Imaging for Label-Free Analysis of Biomolecular Recognition Reactions on Glycolipid Domains. Analytical Chemistry 2012, 84, 6538–6545.

(54) Kirichuk, O.; Srimasorn, S.; Zhang, X.; Roberts, A. R. E.; Coche-Guerente, L.; Kwok, J. C. F.; Bureau, L.; Débarre, D.; Richter, R. P. Competitive Specific Anchorage of Molecules onto Surfaces: Quantitative Control of Grafting Densities and Contamination by Free Anchors. Langmuir 2023, 39, 18410–18423.

(55) Zwang, T. J.; Fletcher, W. R.; Lane, T. J.; Johal, M. S. Quantification of the Layer of Hydration of a Supported Lipid Bilayer. Langmuir 2010, 26, 4598–4601.

(56) Parkkila, P.; Elderdfi, M.; Bunker, A.; Viitala, T. Biophysical Characterization of Supported Lipid Bilayers Using Parallel Dual-Wavelength Surface Plasmon Resonance and Quartz Crystal Microbalance Measurements. Langmuir 2018, 34, 8081–8091.

(57) Horton, M. R.; Reich, C.; Gast, A. P.; Rädler, J. O.; Nickel, B. Structure and Dynamics of Crystalline Protein Layers Bound to Supported Lipid Bilayers. Langmuir : the ACS journal of surfaces and colloids 2007, 23, 6263–6269.

(58) Frey, W.; Schief, W. R.; Vogel, V. Two-Dimensional Crystallization of Streptavidin Studied by Quantitative Brewster Angle Microscopy. Langmuir 1996, 12, 1312–1320.

