## Supplementary Information for "Optical sectioning for reflection interference microscopy: quantitative imaging at soft interfaces"

### Supporting information

**S1 Fig. Setting up the patterned illumination** Detailed procedure to align the lenses and DMD in the illumination path of the microscope. It assumes that the reader starts from a RIC microscope with a transmission illumination arm. Usually, the field diaphragm (II) and sometimes the aperture diaphragm (III and IV) are already included in the optical

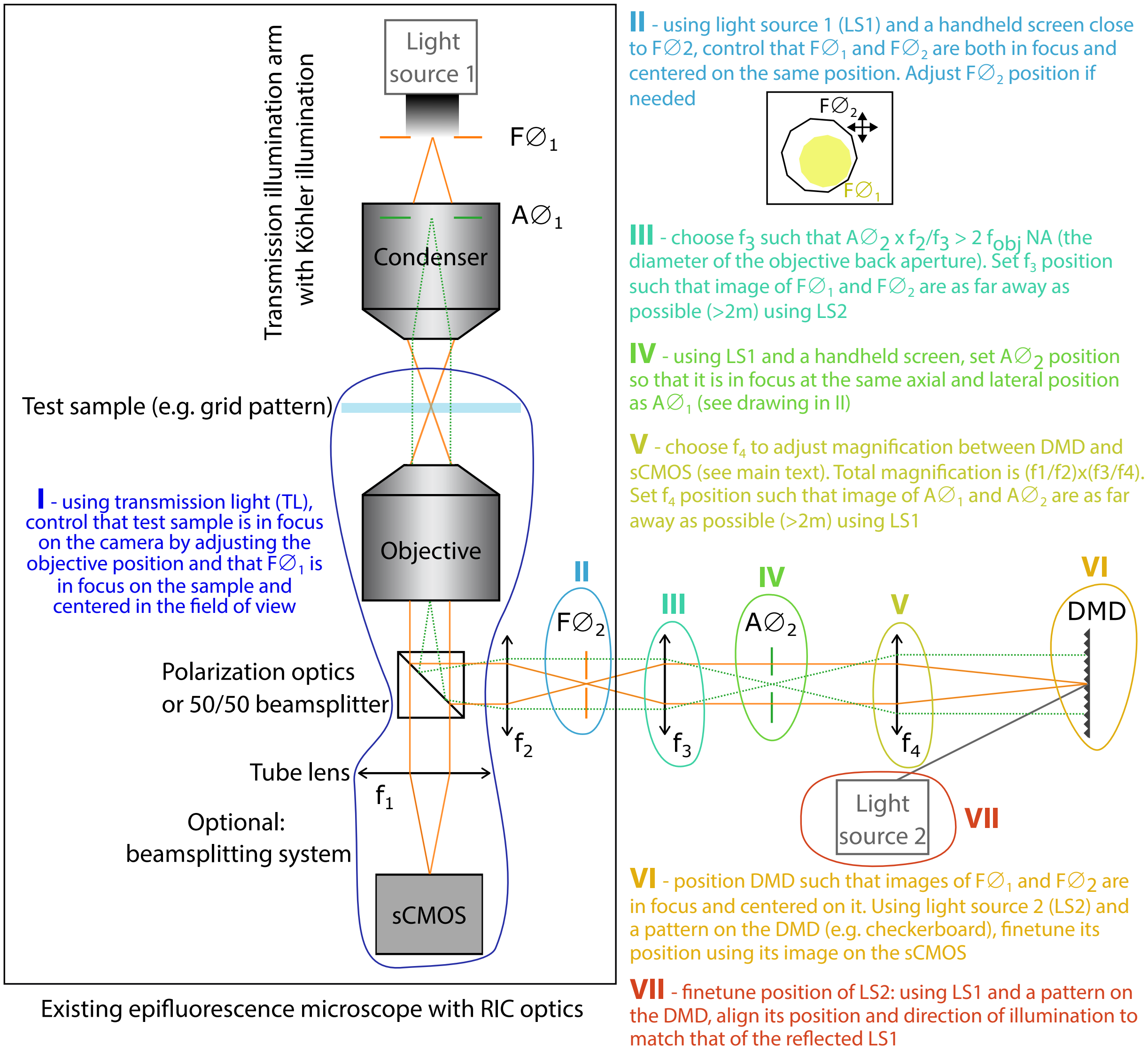


setup, and can be used without further change, leaving only steps I and V-VII to complete. Light path between image planes is shown in orange (sample, field diaphragms, DMD), and between conjugated Fourier plane in light blue (back aperture of the objective, aperture diaphragms). In our system, the DMD pixel size is 6.6$\mu$m while the sCMOS pixel size is 6.5$\mu$m. The DMD axes are chosen parallel to that of the cameras, so that the individual mirrors rotate at a 45 degree angle with the horizontal plane : light source 2 is therefore mounted outside of the horizontal plane. The reader should refer to the details of their DMD to assess the most practical orientation of their device in their optical setup. To adjust the magnification between the DMD and the sCMOS, the focal length of the different lenses are in our case: $f_1$=180mm, $f_2$=75mm, $f_3$=100mm, and $f_4$=200mm. We strongly recommend the use of tube lenses for lenses added to the system outside of the microscope already in place, as they combined low chromatism with low field aberrations, thereby ensuring the maximal image quality over an extended field of view. In our case, $f_3$ and $f_4$ are tube lenses purchased from Thorlabs (TTL-100-A and TTL-200-A, respectively).

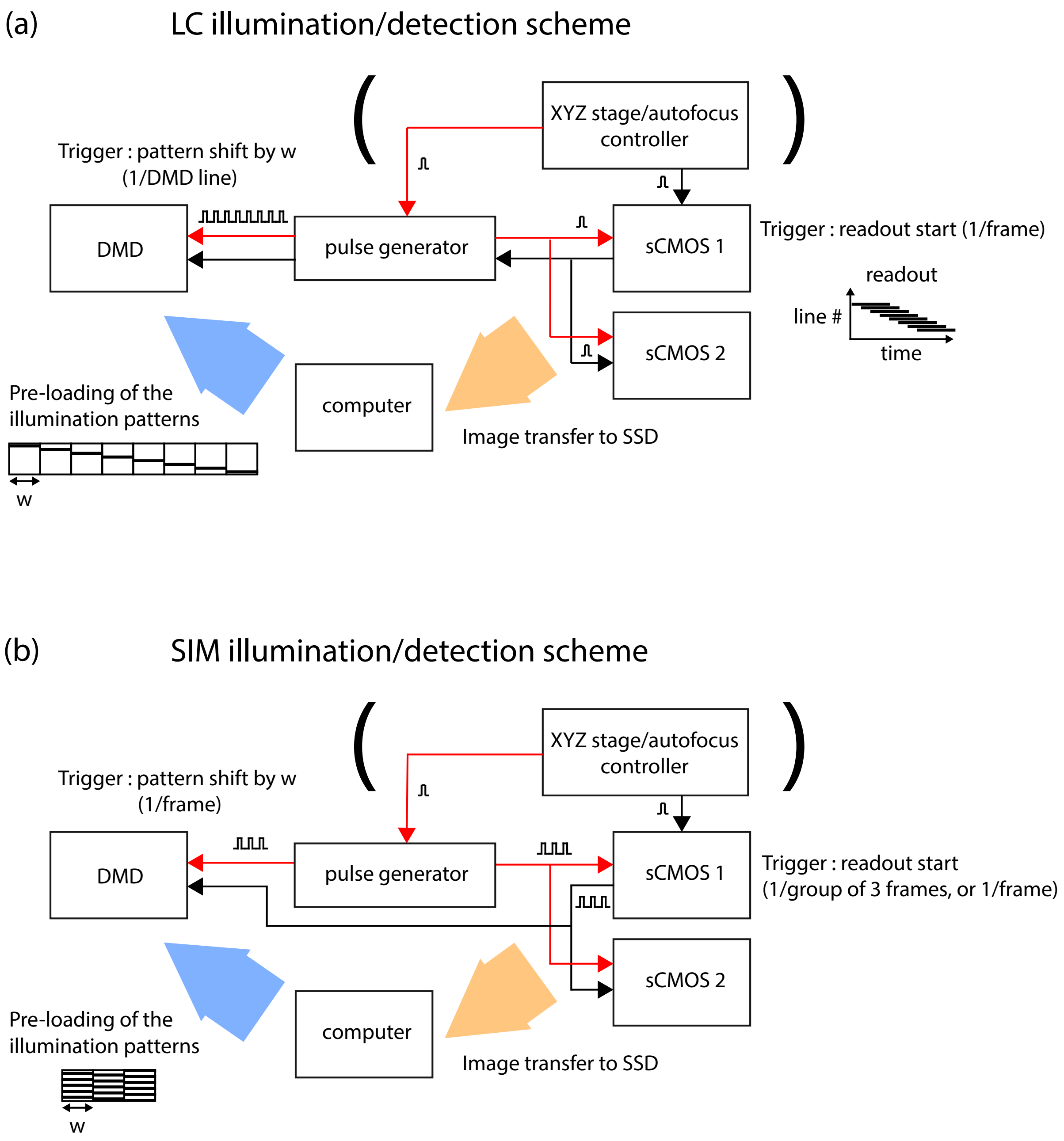


**S2 Fig. Synchronizing light patterning and camera acquisition.** The scheme of hardware synchronization and triggering is presented for LC and SIM. All the instruments are synchronized by TTL pulses using BNC cables for the connections, which makes the control system particularly robust and easy to set up. In both acquisition schemes, acquisition can be externally triggered by a trigger from a displacement or autofocus controller (e.g., to perform a z-scan of the sample or a xy mosaic of 2D images, or ensuring focus lock during a time-lapse acquisition). The use of such an external trigger is however optional. In both cases also, the patterns that will be displayed on the DMD are pre-loaded as a single large image to avoid delays while transferring data to the DMD during imaging. Note that this means that a lag time exists when changing the patterns to be projected, e.g. to change the line width in LC or pattern spatial period in SIM. To achieve good temporal performance, and in particular for SIM real time reconstruction, it is essential to transfer the sCMOS data onto a SSD on the computer, rather than a hard drive.

(a) For LC imaging, the image loaded on the DMD has the dimensions of the DMD (in pixels) times the number of line patterns to be projected. The readout on the cameras must be configured with an adjustable delay for the readout start of each line (”light sheet mode”, for the sCMOS used in this work). This delay matches the delay between the projection of two successive frames on the DMD, divided by the magnification factor between the DMD and the sCMOS. For example, in our setup, one line of pixels on the DMD corresponds to a line of two pixels in thickness on the sCMOS, such that the line rate on the camera must be twice the frame rate on the DMD. Because the refresh time of our DMD is 44 $\mu$s, the time between two camera lines must be at least $\approx$25 $\mu$s and the frame rate is limited to $\approx$40Hz for a $2048 \times 1024$ images. The exposure time is also limited to ensure that at each time point, the readout line width on the camera matches the illumination line width. Typically, the exposure time is set as the line delay ($\geq$ 25 $\mu$s) times the width of the DMD illumination line (in pixels), times the magnification between the DMD and the sCMOS, in pixels (2, in our case). For a line of thickness $\approx$ 1 $\mu$m at the focus of a 60$\times$ objective, this corresponds to an exposure time of 250 $\mu$s.

For the synchronization of the different devices, the master TTL trigger can originate either from one of cameras itself (black path), or from the pulse generator used to synchronize the cameras with the DMD (red path). If it stems from one of the cameras, a single pulse is produced upon the start of the frame acquisition, which directly triggers simultaneous acquisition by the second camera. The same pulse also triggers a series of TTL pulses from the waveform generator that each shift the large image loaded onto the DMD by the DMD width (in pixels), to progressively shift the illumination line. The delay between each pulse is the delay between two successive images displayed on the DMD, and the number of pulses is the total number of images to display to shift the illumination line across the whole field of view. Alternatively, the waveform generator can simultaneously produce a single TTL pulse to trigger the cameras, and a series of pulses to trigger the DMD. This is useful to ensure a reliable acquisition rate on the cameras.

(b) For SIM imaging, except if used to control the frame rate on the camera, the waveform generator is not required. An image composed of the three grid patterns with 1/3 pattern shift is pre-loaded on the DMD, and shifted for each image acquisition. The camera can be in free acquisition mode and output a TTL pulse at the start of each frame readout, or be triggered by an external device. In the second case, our camera can produce internally a burst of three TTL pulses that can be used to acquire the 3 images required for SIM with a minimal delay ("master pulse"). If this feature is not available, a burst of three TTL pulses can be produced by the waveform generator upon triggering by the external device.

Additional details on the optical setup and electronic control are specific to the hardware used here, and are thus of little general interest. However, they are available upon request for the interested reader.

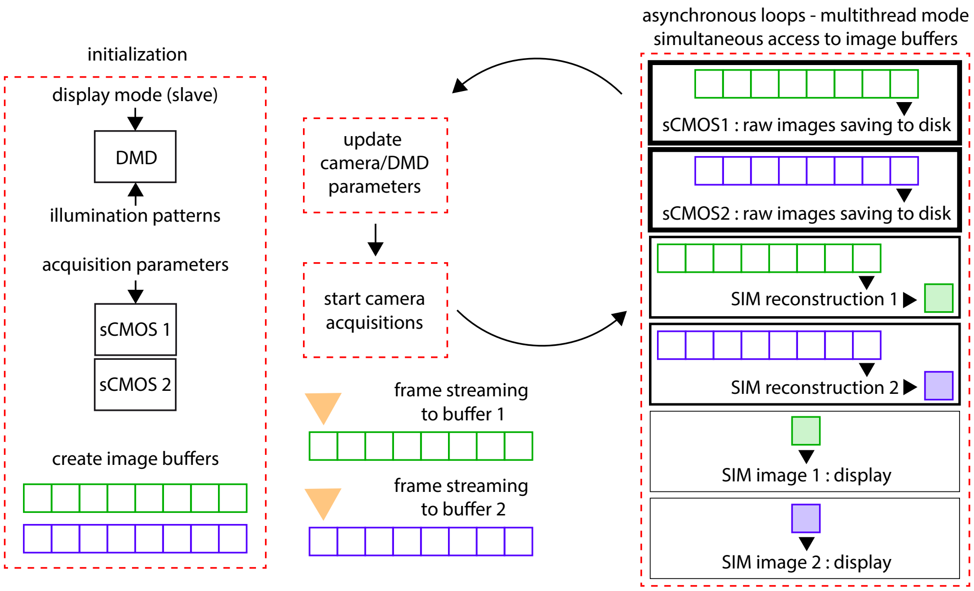


**S3 Fig. Real-time acquisition and processing of SIM images on two sCMOS cameras.** Real-time visualization is essential to use SIM in actual experimental conditions. To this aim, we have developed a home-written software in LabView allowing simultaneous image acquisition, saving, reconstruction and display for two 2048 × 2048 pixel cameras running in 16 bits. The program operates at up to 20Hz (single camera) or 10Hz (two cameras) on a 8 cores, 64Mo-RAM acquisition computer from 2017 running under windows 10 and equipped with a 128 Go SSD. These requirements are largely surpassed by any recent computer, onto which the program should run with even better performance. To achieve real-time performance, two elements are key: first, one has to take advantage of the multiple cores by running parallel operations (native under LabView but challenging e.g. in Python at present). Secondly, the different operations (streaming from the camera to the RAM, saving to the solid-state drive, SIM reconstruction, and display) must be performed independently, so that waiting times between the different parts of the program are eliminated. To this aim, we used asynchronous loops set to run each on a different processor, and store the streamed images on two buffers in the RAM allowing simultaneous access and writing. Finally, the different loops are assigned different priorities: saving has the highest priority so as to ensure that no raw frames are missed for subsequent analysis (a flag indicates if this occurs during saving). We choose to save the raw images here, so that alternative, slower reconstruction algorithms can be used offline if needed instead of the canonical 3-frames standard deviation method that is used for real-time display. SIM reconstruction is next, and display has the lowest priority as an occasional missed frame at relatively high frame rate does not affect the user much and does not impact subsequent analysis of the saved data. To improve the frame rate and facilitate experimental use (e.g., focussing, lateral displacement, etc.), we use a rolling SIM reconstruction similar to the rolling average commonly available on commercial acquisition software: for reconstruction, the last three images in the image buffer are used to compute their standard deviation (corresponding to the SIM image), such that these three images each provide one of the three phases for the SIM pattern, but that they are only shifted by one image (or time step) between two reconstructions. This provide a reconstruction at the acquisition frame rate (instead of one third of it), with a 3-frame smoothing effect. Additional details on the program are specific to the hardware and LabView version used here, and are thus of little general interest. Any information about our acquisition program is nevertheless available upon request to help the interested reader set up their own acquisition program.

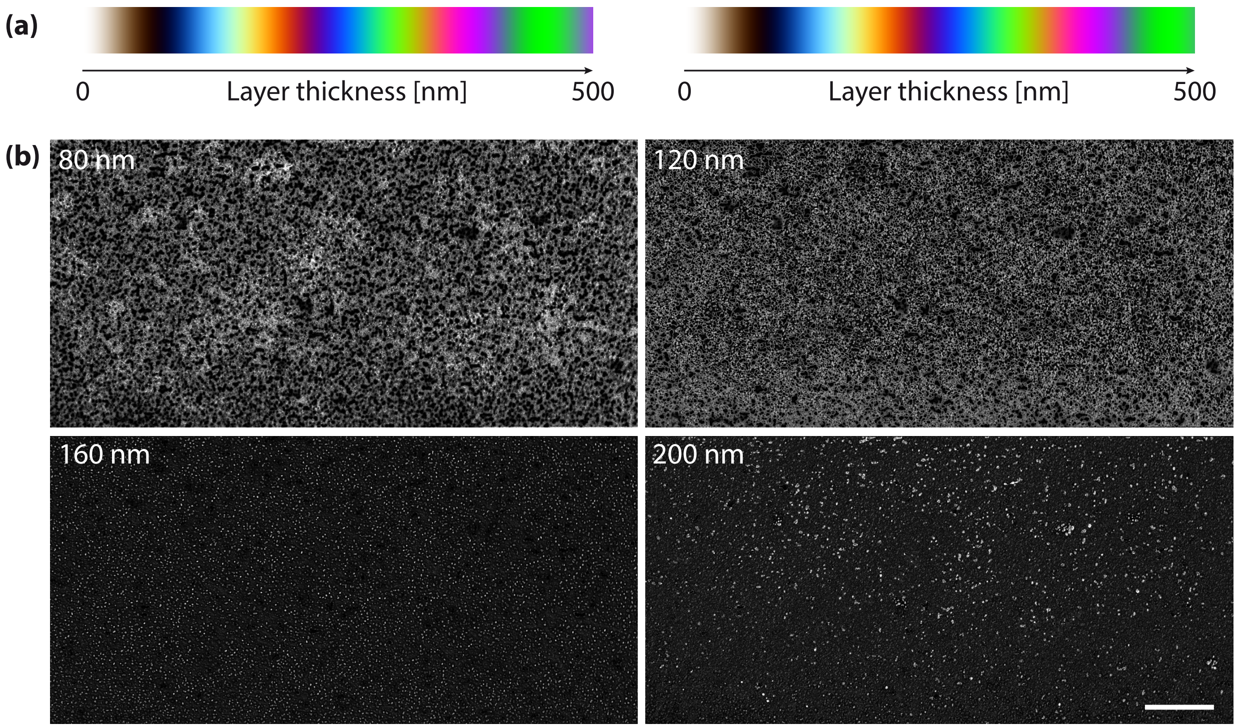


**S4 Fig. Classification of colour RICM image from Fig.4a.** (a) Colour scales calculated for a high density film (n=1.47, left) or a 50% water content (n=1.4, right), showing a very weak dependence of the colour scale to the exact film hydration. (b) Four examples of probability maps corresponding to heights 70-90nm, 110-130nm, 150-170nm, and 190-210nm, obtained with the Weka Trainable Segmentation plugin. The probabilities scale between 0 (black) and 1 (white). Scale bar, 100 $\mu$m.

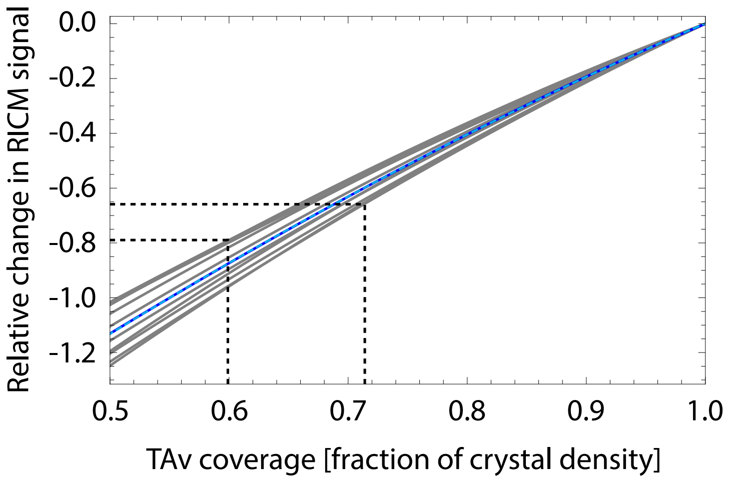


**S5 Fig. Calculated optical response to a change in the TAv surface density.** The relative change in RICM intensity with respect to the case of a crystalline monolayer of TAv is plotted against the density of TAv normalized to that of 2D crystals. The scenario exposed in the methods section yields the blue curve, fitted with a parabola (cyan dotted curve). Other curves correspond to variations of the model parameters, based on experimental uncertainties in the various existing studies, to estimate its robustness: $\pm 0.02$ for TAv crystal refractive index, $\pm 0.01$ for the DOPC refractive index, $\pm 0.2$ nm in the SLB thickness, $\pm 0.5$nm in TAv layer thickness, $\pm 0.2$nm in water layer thickness. Accounting for the uncertainty in the model increases the uncertainty on the calculated TAv density to approximately $\pm 10\%$.

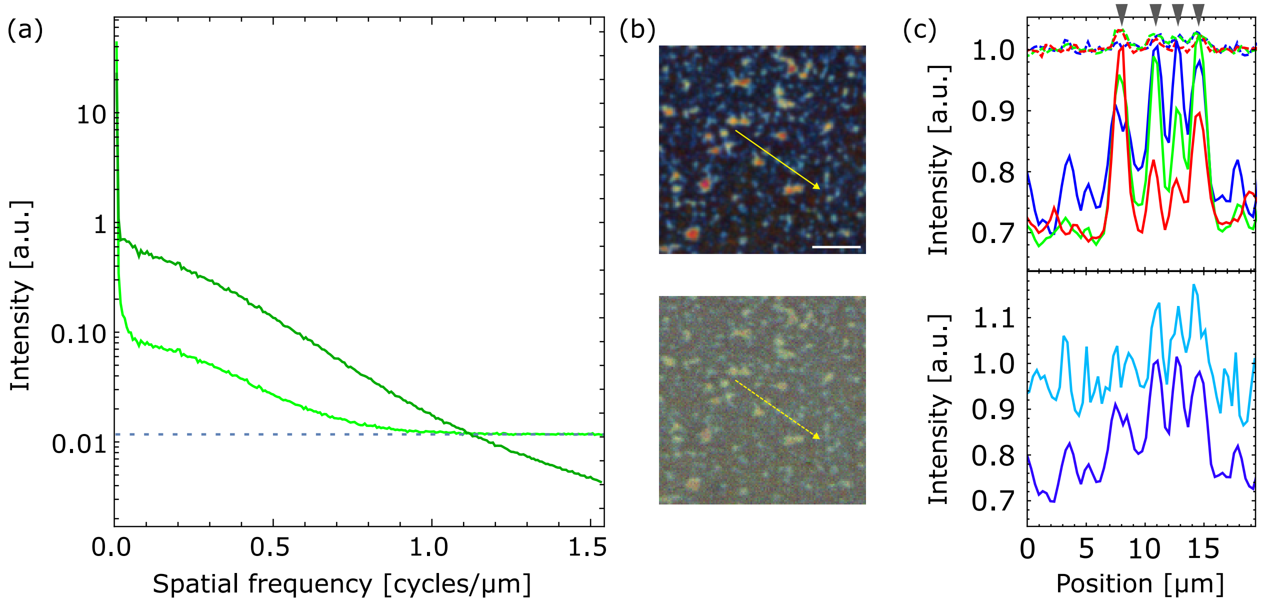


**S6 Fig. Additional quantifications on RICM images from Fig.4a.** (a) Modulation frequency spectra from the green channel of the LC (dark green) and WF (light green) RIC images. The frequency cutoff in the absence of noise is calculated to be around 2 INA/$\lambda$=3 cycles/$\mu$m, while sampling by the camera limits detected frequencies to 1.5 cycles/$\mu$m. Useful frequency content is detected in LC RIC images up to this limit, whereas WF RIC image frequency spectrum is dominated by noise above about 1 cycles/$\mu$m (blue dashed line).(b), detail of images from Fig. 4(a) displaying the line over which profiles in (c) are measured. Top, LC RICM; bottom, WF RICM. Scale bar, 10 $\mu$m. (c), top, corresponding profiles for LC (solid lines) and WF (dashed lines) at 453 (blue), 532 (green) and 632 nm (red). The contrast of signal variations are of much larger amplitude in the LC image. Bottom, to better compare the profiles, WF profile at 453nm (light blue) is drawn with increased contrast to match that of the LC profile (dark blue). The offset of the curves are arbitrary. Arrowheads on top of the graph indicate the 4 structures along the yellow arrow that are clearly visible in the original image.

**S1 Video. Monitoring of TAv crystallization over time.** Sectioned (left) and wide-field (right) images of a biotinylated SLB incubated with TAv, showing the growth of crystals over time. The time indicated on the movie is the time after the start of incubation. See also Fig.3 and Methods.
